# Evolutionary parallelisms of pectoral and pelvic network-anatomy from fins to limbs

**DOI:** 10.1101/374504

**Authors:** Borja Esteve-Altava, Stephanie E. Pierce, Julia L. Molnar, Peter Johnston, Rui Diogo, John R. Hutchinson

## Abstract

Pectoral and pelvic lobe-fins transformed into fore- and hindlimbs during the Devonian period, enabling the water-to-land transition in tetrapods. In the timespan of ~60 million years, transitional forms evolved, spanning a wide range of morphologies. Here we traced the evolution of well-articulated appendicular skeletons across the fins-to-limbs transition, using a network-based approach and phylogenetic tools to quantify and compare topological features of skeletal anatomy of fins and limbs. We show that the topological arrangement of bones in the pectoral and pelvic appendages evolved in parallel during the fins-to-limbs transition, occupying overlapping regions of the morphospace, following a directional mode of evolution, and decreasing their disparity over time. We identify the presence of digits as the morphological novelty triggering significant topological changes that clearly discriminated limbs from fins. The origin of digits caused an evolutionary shift towards appendages that were less densely and heterogeneously connected, but more assortative and modular. Topological disparity likewise decreased for both appendages: for the pectoral appendage, until the origin of amniotes; for the pelvic appendage, until a time concomitant with the earliest-known tetrapod tracks. Finally, we tested and rejected the presence of a pectoral-pelvic similarity bottleneck for the network-anatomy of appendages at the origin of tetrapods. We interpret our findings in the context of a dynamic compromise between possibly different functional demands in pectoral and pelvic appendages during the water-to-land transition and a shared developmental program constraining the evolvability of limbs.

---

The evolution of tetrapod limbs from fish fins is heralded as one of the most important vertebrate morphological and functional transitions^1–8^. Establishing what makes an appendage a fin or a limb is key to properly characterizing the fins-to-limbs transition^3^. Functional criteria are of limited use because of the general consensus that limbs first evolved to move under water^5,9^. Developmental and palaeontological studies place the distinction between fins and limbs in the most distal region, which bears the carpals/tarsals and digits in limbs and the radials and dermal lepidotrichia in fins^3,10^. The distinction between fins and limbs blurs when we look at the lobe-fins of transitional tetrapodomorphs, such as *Eusthenopteron*, *Gogonasus*, and *Tiktaalik*^11–13^. Both lobe-fins and limbs share a division of the appendicular skeleton into three endoskeletal domains^14^, of which the most distal one shows the greatest differentiation between sarcopterygian fishes (i.e., multi-patterned radial bones) and tetrapods (i.e., autopod with a mesopod and digits). Although in the past, researchers have disagreed about whether a zeugopod-mesopod boundary (wrist/ankle)^15,16^ or the presence of digits alone^3^ is sufficient to define limbs, the current general convention is to define “true” limbs as appendages with digits^6^. Even though the anatomical organization/topology of the distal radials and the autopod superficially look similar (i.e., a series of skeletal elements joined proximodistally)^3,17^ and they share a common genetic control or “deep homology”^18^, their anatomical similarity has never been assessed quantitatively. Moreover, pectoral and pelvic lobe-fins evolved into limbs in tandem during the fins-to-limbs transition, made possible due to the recruitment of a common developmental genetic toolkit^2,19^. Because pectoral and pelvic appendages were originally different in their anatomy^20^—and still are regarding the genetics of girdle development^21^—we would expect to see a mix of evolutionary parallelisms/convergences (homoplasy) and divergences, as shared and specific developmental programs and biomechanical functions intertwined with each other during the fins-to-limb transition. Such a mix might result from compromises between these “evo-devo” and “evo-biomechanical” constraints.

As appendages evolved, the anatomical similarity between pectoral and pelvic appendages also evolved. Various authors have proposed alternative bottlenecks during evolution for the pectoral-pelvic similarity (reviewed in refs ^7,22^); these evolutionary bottlenecks represent times when pectoral and pelvic appendages showed a greater anatomical similarity to each other (i.e., their morphologies showed less disparity). Based on skeletal and muscular anatomical and developmental features^20, 23–25^, pectoral-pelvic similarity bottlenecks have been proposed for the origins of ray-finned fishes, coelacanths, tetrapodomorphs, and tetrapods. In a recent study comparing the musculoskeletal network-anatomy of whole appendages^22^, we found evidence for a pectoral-pelvic similarity bottleneck at the origin of sarcopterygians (as proposed by refs ^3,7,20,25^); but not at the origin of tetrapods (as these same studies proposed). However, our previous work focused only on the network-anatomy of extant taxa^22^. To further test the presence of a pectoral-pelvic similarity bottleneck at the origin of tetrapods for the network-anatomy of the skeleton, we have analysed a broader sample of extinct sarcopterygian fishes and early tetrapods across the fins-to-limbs transition, including *Sauripterus*, *Eusthenopteron*, *Gogonasus*, *Tiktaalik*, *Acanthostega*, *Ichthyostega*, *Tulerpeton*, *Balanerpeton*, *Eryops*, *Seymouria*, and *Westlothiana*; for which the fully-articulated pectoral and/or pelvic anatomy is reasonably well known. A prediction of the hypothesis of a pectoral-pelvic similarity bottleneck at the origin of tetrapods is that taxa closer to the split of Tetrapoda within sarcopterygians—where the bottleneck is—will have a greater pectoral-pelvic similarity (or lower disparity) than taxa that are farther away from the bottleneck.

To compare the skeletal anatomy of appendages in extinct and extant forms across the fins-to-limbs transition, and better characterize anatomical parallelism/convergence and divergence between pectoral and pelvic appendages, we focused our analysis on their anatomical organization or network-anatomy; that is, the topological arrangement/pattern of skeletal elements of fully-articulated appendages (**Fig. 1a**). This level of abstraction allowed us to compare evolutionary changes that are not amenable to quantification using other morphometric methods due to the large disparity of forms and presence/absence of parts between pectoral and pelvic fins and limbs^22,26,27^. Furthermore, the abstraction retains biological meaning in that the contacts between skeletal elements reflect potential direct developmental and biomechanical interactions; for example, ontogenetic sequences of ossification or embryonic interaction, and joint reaction forces or ranges of motion. Using a network-based approach^22,28,29^, we modelled the skeleton of fully-articulated appendages as networks, in which nodes code for bones and links code for physical contact in a standardized resting pose. We compared the evolution of eight network-based topological variables (see *Methods* for details) in a phylogenetic context (**Fig. 1b**) to test whether (1) there are topological differences between fins and limbs and between pectoral and pelvic appendages, (2) pectoral and pelvic anatomy followed convergent/parallel or divergent modes of evolution during the fins-to-limbs transition, and (3) there was an evolutionary bottleneck in pectoral-pelvic similarity in tetrapods. We tested these hypotheses by comparing the occupation of appendicular morphospace, estimating shift of evolutionary regimes, describing the evolution of disparity through time, and testing bottlenecks with phylogenetic regressions.

**Figure 1.**
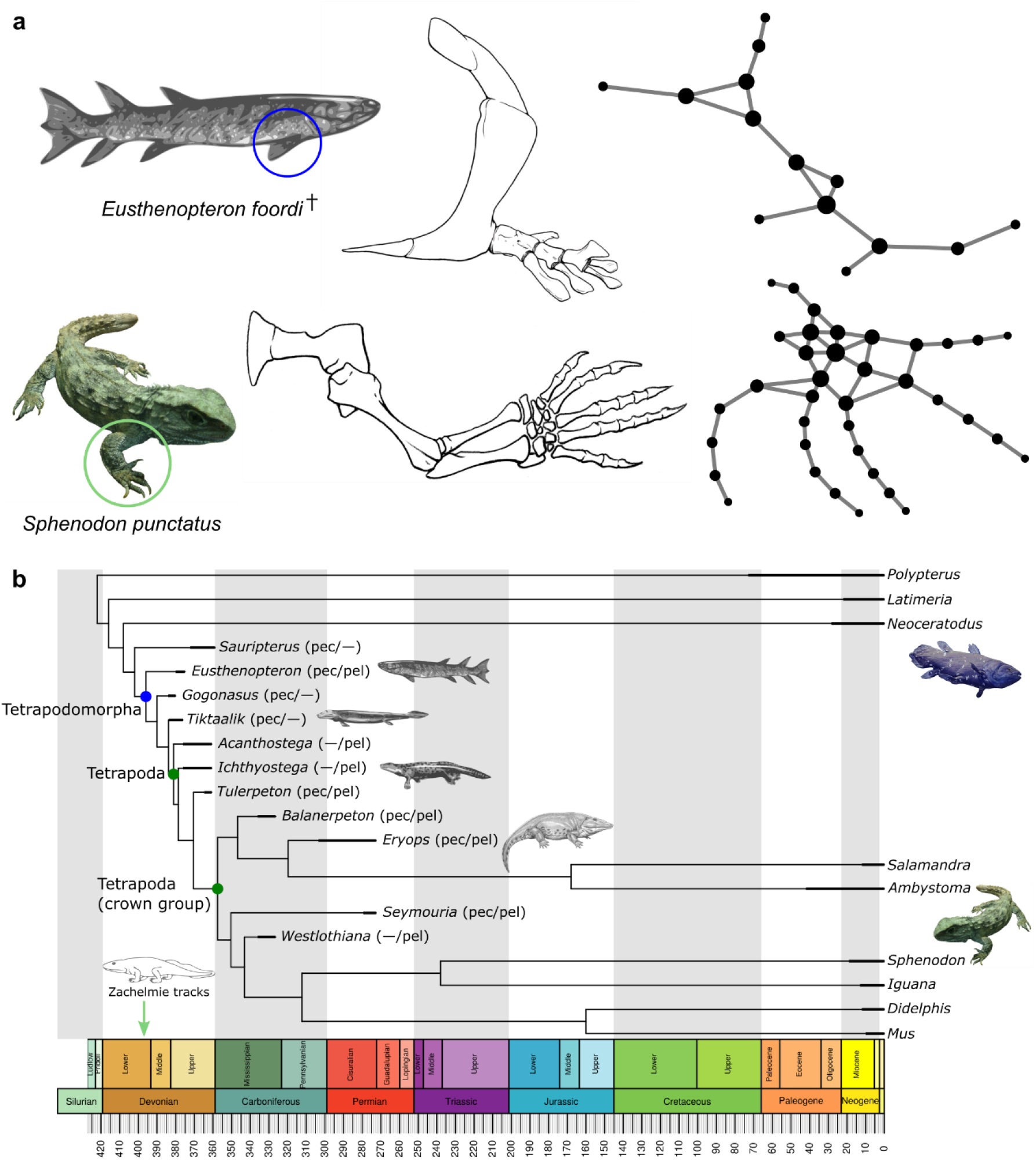
Network abstraction and phylogenetic context of the study. **a**, Representative network models of the skeletal anatomy of a fin and a limb; network nodes represent the bones of the appendage and the links connecting them represent their physical articulations or joints. Note that anatomical-network models are purely topological; thus, information about the size, shape, and positioning of bones is not part of the model. Node size is drawn proportional to the bone’s number of articulations. **b**, Time calibrated phylogenetic tree assembled for this study showing which taxa have complete information for pectoral and pelvic appendages: pec/pel, both appendages found in articulation or completely reconstructed; –, complete appendage not preserved. Image of tetrapod outline by Mateus Zica (GFDL), body fossil restorations by Nobu Tamura (CC BY-SA 3.0), coelacanth by Zoo Firma (CC BY-SA 3.0), and tuatara by Tim Vickers (CC BY-SA 3.0).

## RESULTS

### Topological Discrimination of Appendages

The network-anatomy of the appendicular skeleton varied for each taxon and between pectoral and pelvic regions (**Table 1**). We used a Principal Component Analysis (PCA) to visualize global patterns of topological variance across anatomical networks and test for differences between fins and limbs, pectoral and pelvic appendages, and extinct and extant species.

**Table 1.**
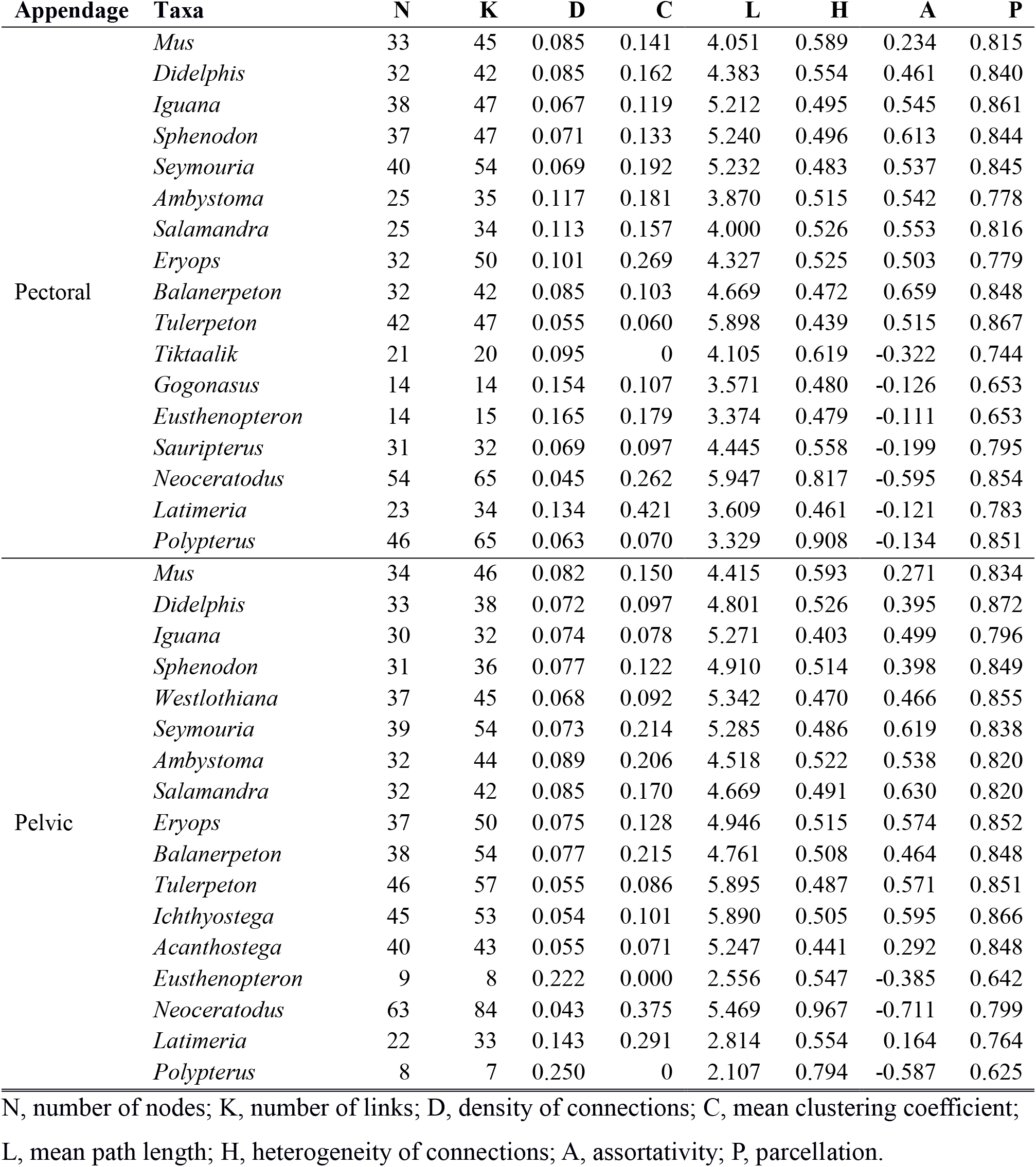
Values of network variables measured on each network model.

The first two PCA components explained 81.8% of the total topological variation among appendages (**Fig. 2a**). The first axis of variation (58.9%) broadly discriminated between (*i*) more modular (higher P) sparsely connected (lower D) appendages, such as limbs, and (*ii*) less modular (lower P) densely packed (higher D) appendages, such as lobe-fins. The second axis of variation (22.9%) broadly discriminated between (*i*) more regular appendages, in which bones tend to have the same number of articulations (lower H) and which contact bones with a similar number of articulations (high A), such as limbs and anatomically plesiomorphic lobe-fins, and (*ii*) more heterogeneous appendages, in which bones have a varying number of articulations (higher H) and which preferentially contact bones with a different number of articulations (lower A), such as in the anatomically derived lobe-fins of *Neoceratodus*.

**Figure 2.**
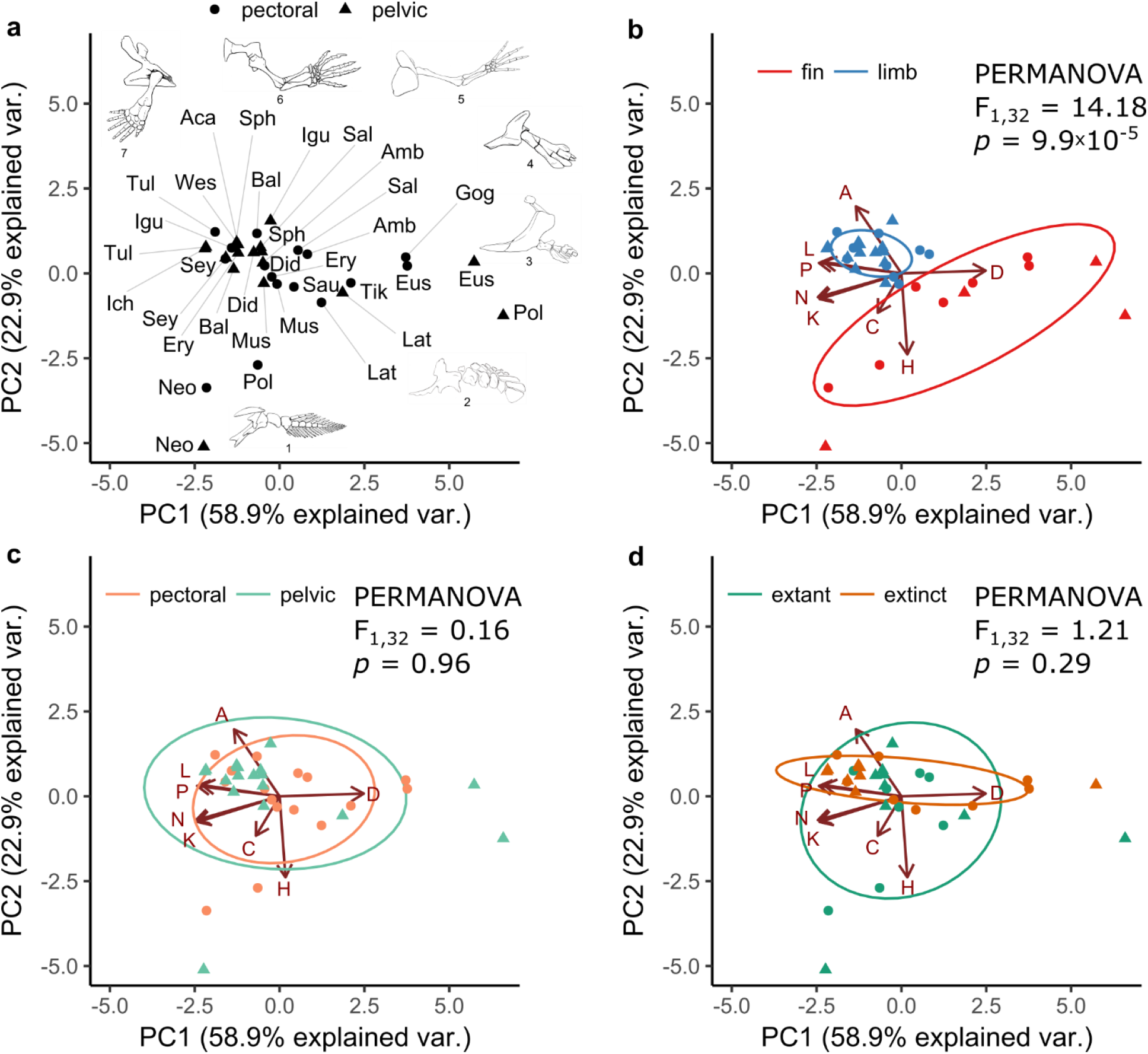
Biplots of the first two PCA components of topological variables for pectoral (circles) and pelvic (triangles) appendages combined. **a,** Distribution of pectoral and pelvic appendages for each taxon in the sample (abbreviated by the first three letters of the genera). For reference we included line drawings representing different appendages: (1) *Neoceratodus* pectoral, (2) *Latimeria* pelvic, (3) *Eusthenopteron* pelvic, (4) *Gogonasus* pectoral, (5) *Salamandra* pectoral, (6) *Acanthostega* pelvic, and (7) *Sphenodon* pectoral. **b**, Comparison of limbs *vs.* fins (i.e., appendages with and without digits, respectively) showed that they occupy different regions of the morphospace. **c**, Comparison of pectoral *vs.* pelvic appendages showed that they occupy overlapping regions of the morphospace. **d**, Comparison of extant *vs.* extinct taxa showed that they occupy the PC2 axis in different ways. Red arrows show the contribution to the first two PCA components of each network variable: (N) nodes, (K) links, (D) density, (C) clustering coefficient, (L) path length, (H) heterogeneity, (A) assortativity, and (P) parcellation.

A permutational multivariate analysis of variance (PERMANOVA) showed a statistically significant difference in topological variability between fins and limbs (F_1,32_ = 14.18, *p* = 9.9×10^−5^; **Fig. 2b**). The assortativity of appendages (i.e., tendency of bones to contact bones with a similar number of connections) was the main discriminator between fins and limbs, which occupied opposite positions along the assortativity*-*axis of variation. Limbs had larger and positive values, whereas fins had lower and negative values. We can explain this difference by the presence of the autopod in limbs and, more specifically, by the presence of digits. Phalanges and, to a lesser extent, carpal/tarsal bones, tend to articulate with other autopodial bones with a similar number of articulations. For example, phalanges connect in a proximodistal series to other phalanges or metacarpals/metatarsals, so that most phalanges have two articulations (one proximal one distal) to other phalanges that also have two articulations (hence A increases). A similar pattern may occur among carpal/tarsal bones because of their nearly-polygonal shapes. PERMANOVA showed no significant difference in topological variability between pectoral and pelvic appendages (F_1,32_ = 0.16, *p* = 0.96; **Fig. 2c**) and between extinct and extant species (F_1,32_ = 1.21, *p* = 0.29; **Fig. 2d**). Pectoral and pelvic appendages did not occupy different areas of the morphospace, which indicates that they share a similar topological organization. Although extinct and extant taxa are statistically indistinguishable, they appear to occupy slightly different areas of morphospace differently: extant species varied equally along PC1 and PC2 axes, while the variance in extinct species is primarily concentrated along the PC1 axis and barely varied along the PC2 axis.

### Evolution of Pectoral and Pelvic Appendages

We assessed the potential for parallel/convergent and divergent changes in topology for pectoral and pelvic appendages by estimating shifts in evolutionary regimes (SURFACE) and analysing disparity through time (DTT).

The SURFACE analysis on PC1 and PC2 estimated a shift in mean values at the root branch of Tetrapoda for the complete sample of pectoral appendages (**Fig. 3a**); thus, dividing the sample into two regimes, one for radial-bearing taxa and another for digit-bearing taxa. The signal-to-noise ratio of this estimated pattern was higher than one (PC1 = 2.03; PC2 = 15.87), which indicates a high effect size of both variables in discriminating groups and adequate power to detect shifts. Comparisons of alternative evolutionary models using AIC weights showed that an Ornstein-Uhlenbeck model with multiple rates of change (sigma) and optimal means (theta) best explains the evolution of pectoral appendages (**Supplementary Table 1**). A phylogenetic MANOVA confirmed a statistically significant difference between pectoral fins and pectoral limbs (F_1,32_ = 25.87, *p* = 9.9×10^−5^). A similar pattern was found when we analysed only those taxa for which we had a pelvic correspondence in the sample. In this pectoral subsample, the estimated new regime also included *Eusthenopteron* (**Fig. 3b**), which placed the shift in mean values at the root branch of Tetrapodomorpha (rather than Tetrapoda). The signal-to-noise ratio for the pectoral subsample was above one (PC1 = 1.76; PC2 = 1.73); enough to detect a difference, but weaker than for the complete sample. Likewise, an Ornstein-Uhlenbeck model was the best fit and phylogenetic MANOVA showed a statistically significant difference between the two groups, both including *Eusthenopteron* (F_2,11_ = 19.65, *p* = 0.0052) and excluding it (F_2,11_ = 20.2, *p* = 6.9×10^−4^). The congruence of results added support to the estimated shift in topological organization during the fins-to-limbs transition.

**Figure 3.**
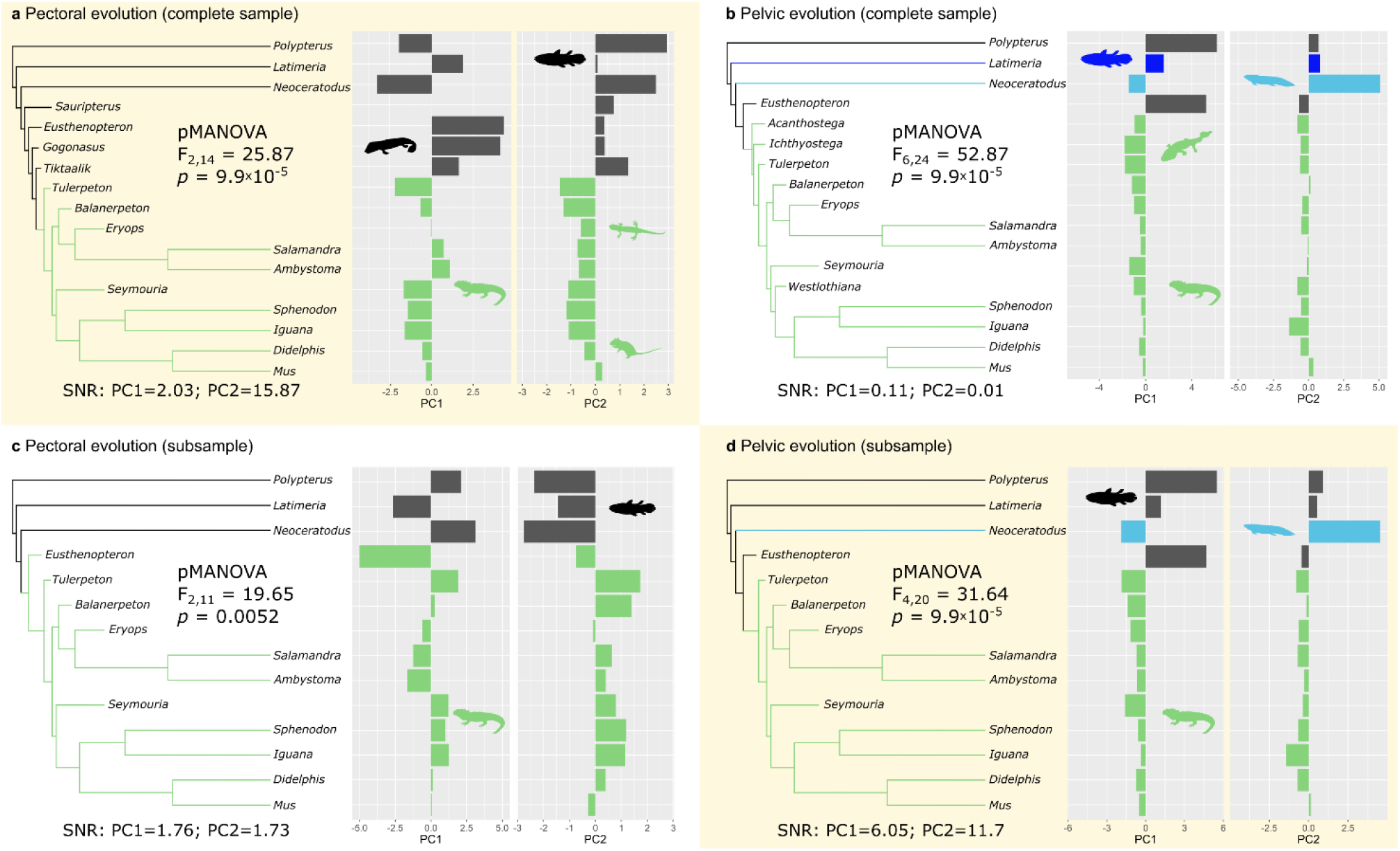
Estimated evolutionary shifts using SURFACE. **a**, Estimation using the complete sample of pectoral appendages. **b**, Estimation using the subsample of pectoral appendages having both appendages represented in the sample. **c**, Estimation using the complete sample of pelvic appendages. **d**, Estimation using the subsample of pelvic appendages having both appendages represented in the sample. Both appendages showed a regime shift in Tetrapoda, indicated in green. Other single-lineage shifts found in lobe-finned fished were highlighted in shades of blue. Yellow background marks SURFACE tests with a signal-to-noise ratio (SNR) above one. Significant SNR indicated evolutionary shifts at the origin of digit-bearing taxa for both pectoral and pelvic appendages, with a potential independent shift in *Neoceratodus* pelvic fin evolution. Evolutionary shifts toward different topologies were validated by phylogenetic MANOVA tests.

The SURFACE analysis for the complete sample of pelvic appendages estimated three shifts of mean values (**Fig. 3c**): one at the root branch of Tetrapoda, and another two in the lineage of *Latimeria* and *Neoceratodus*. However, the signal-to-noise ratio of the estimated pattern was below one for both variables (PC1 = 0.11; PC2 = 0.01), which indicated a lack of effect size and power to discriminate between groups. Comparisons of alternative models using AIC weights showed that an Ornstein-Uhlenbeck model with multiple optimal means (theta) better fitted the evolution of pelvic appendages (**Supplementary Table 1**). Despite the low signal-to-noise ratio, phylogenetic MANOVA confirmed a statistically significant difference between radial-bearing taxa vs. digit-bearing taxa (F_2,14_ = 47.17, *p* = 9.9×10^−5^). A similar pattern was found when we analysed only those taxa for which we have a pectoral correspondence in the sample. In this pelvic subsample, only two shifts were estimated: one at the root branch of Tetrapoda and another in the lineage of *Neoceratodus* (**Fig. 3d**). In contrast to what happened for the complete sample of pectoral appendages, these regime shifts had a signal-to-noise ratio higher than one for both variables (PC1 = 6.05; PC2 = 11.7). Like for the complete sample, an Ornstein-Uhlenbeck model was the best fit and a phylogenetic MANOVA showed a statistically significant difference between radial-bearing taxa and digit-bearing taxa (F_2,11_ = 34.72, *p* = 0002).

Topological disparity decreased through time for pectoral and pelvic appendages alike (**Fig. 4a**; *solid black line*). DTT tests for the complete samples showed a statistically significant decrease of disparity in pectoral (MDI = –0.37, *p* = 1.2×10^−5^) and pelvic appendages (MDI = –0.38, *p* = 8.6×10^−9^) in the timespan between the origins of sarcopterygians and amniotes (**Fig. 4a**; *red dashed lines*). For pectoral appendages, the decay was more exponential between sarcopterygians and amniotes. For pelvic appendages, there was a pronounced linear decay until a turning point that was roughly concurrent with the age of the oldest purported tetrapod tracks (**Fig. 4a**; *green dashed line*); after that point disparity continued mostly steady through time. As the pelvic appendage dataset contains few tetrapodomorph fish, we subsampled the two datasets to contain similar taxonomic spread. The subsampled DTT showed similar patterns between pectoral and pelvic appendages, with a change in disparity coincident with the age of the oldest tetrapod tracks (**Fig. 4b**). This match indicates that removal of tetrapodomorphs fishes (*Sauripterus*, *Gogonasus*, and *Tiktaalik*) reduced the disparity of appendages in the time-frame of interest.

**Figure 4.**
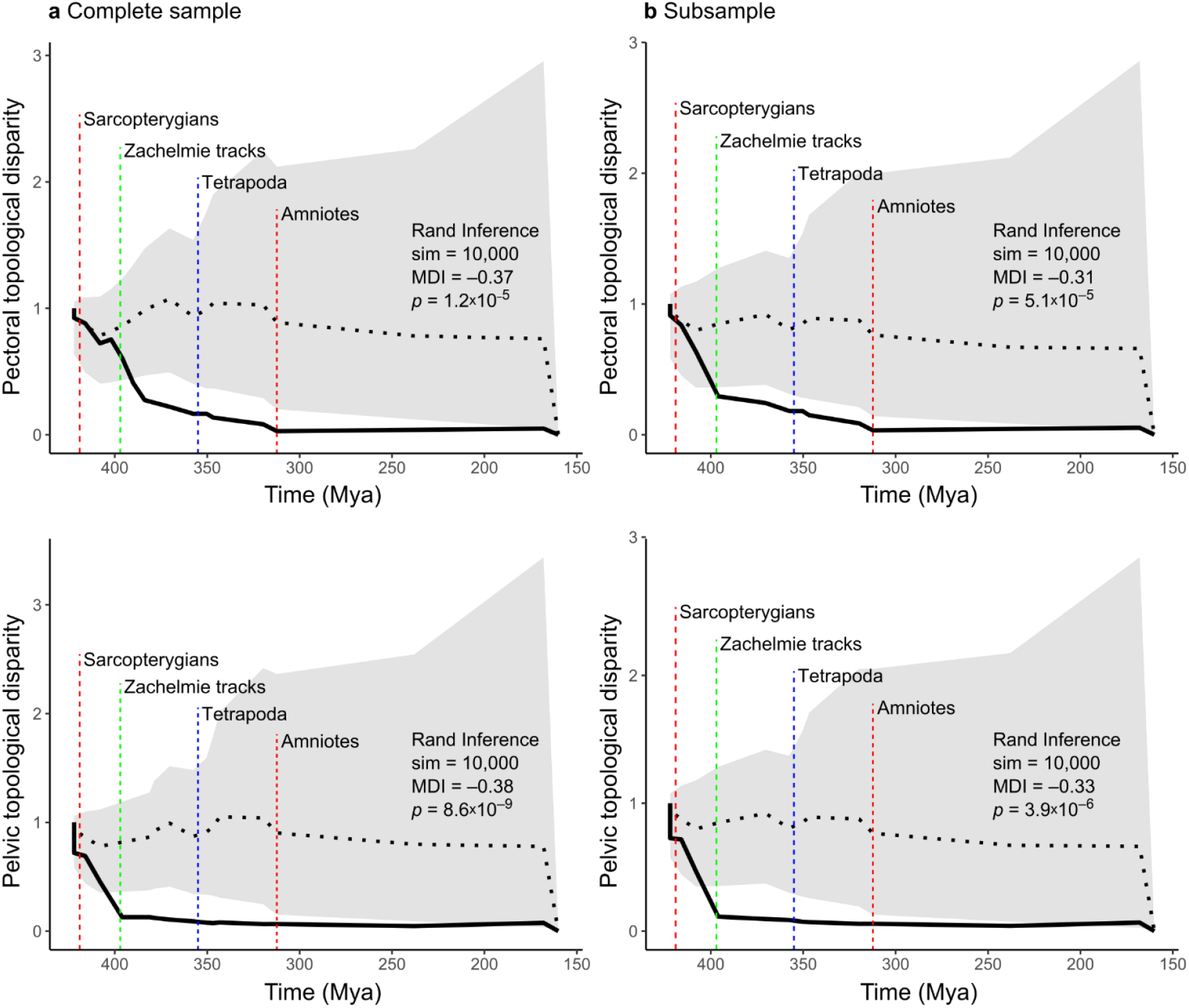
Topological disparity through time for pectoral and pelvic appendages. **a**, Complete samples. **b**, Subsamples with only taxa having both appendages modelled. There was a decay in disparity (*solid black line*) for both appendages, which varied in the rate of decay in different periods of time. Grey areas show the 95% CIs of the expected disparity based on 10,000 phylogenetic simulations under Brownian motion. Horizontal black dotted line shows the mean of disparity for all simulations. Vertical colour dashed lines mark key events (e.g., divergence times) in sarcopterygian evolution for reference.

### Pectoral-Pelvic Similarity Bottleneck

To test the similarity bottleneck hypothesis, we performed a phylogenetic generalized least square (PGLS) regressions of pectoral-pelvic disparity against time from Tetrapoda, for each of the topological variables. As a proxy for similarity we used disparity, calculated as the absolute residual of pectoral and pelvic network variables to the identity line (lower disparity = higher similarity).

The bottleneck hypothesis predicts regression slopes significantly greater than zero for disparity against time. Because the bottleneck marks the point of minimum disparity (maximum similarity), taxa far from the bottleneck would have higher disparity (lower similarity) than taxa close to the bottleneck. None of the PGLS regressions were statistically significant (**Fig. 5**), which means that the topological arrangement of the pectoral-pelvic appendages does not support a similarity bottleneck at the origin of Tetrapoda.

**Figure 5.**
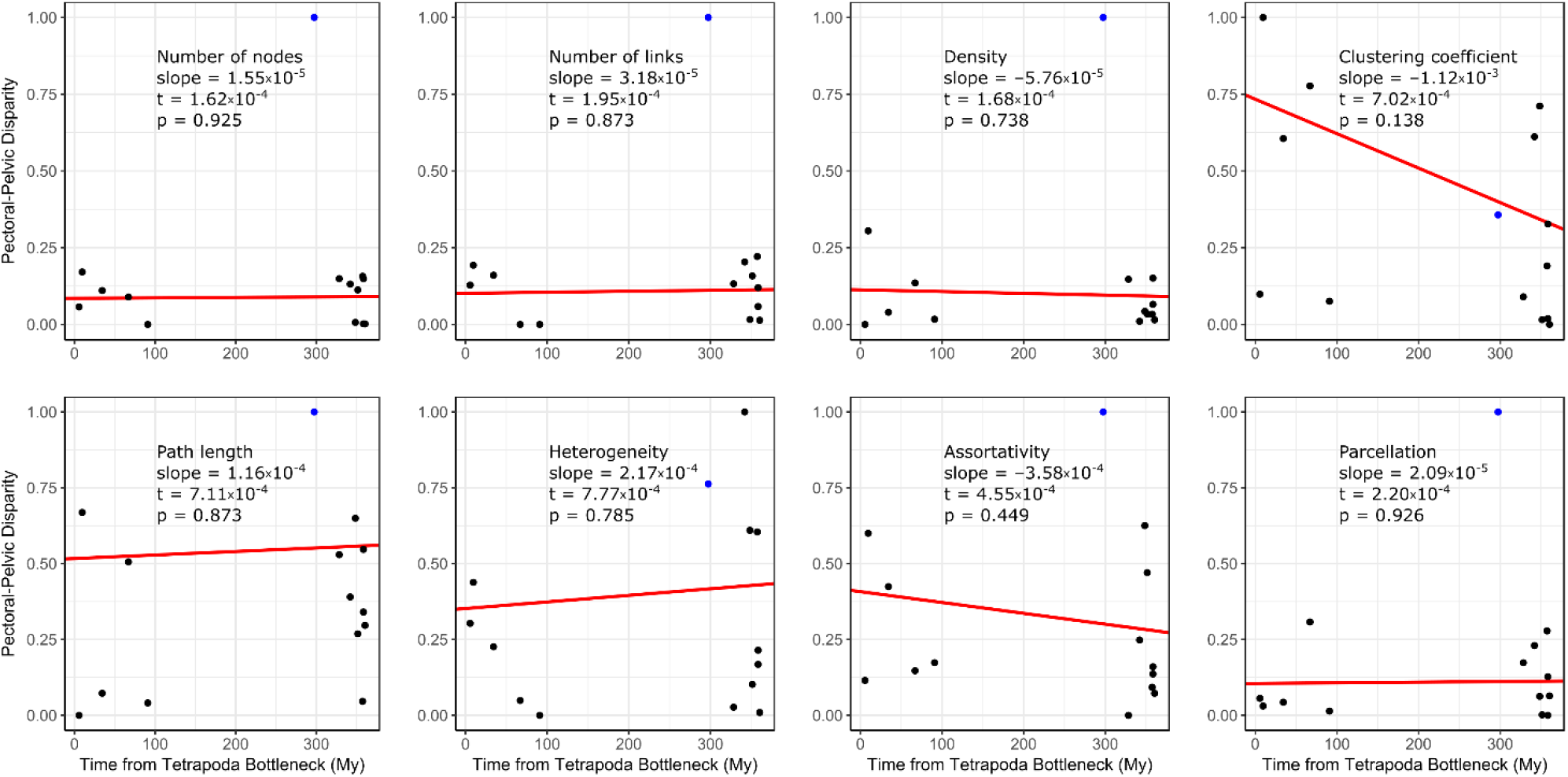
Testing the pectoral-pelvic similarity bottleneck hypothesis for Tetrapoda. PGLS regression slopes (in red) were not different from zero in any of the eight variables analysed; thus, rejecting the presence of a pectoral-pelvic similarity bottleneck for the topological organization of the appendicular skeleton. A significantly positive slope would be consistent with a bottleneck. Blue dot marks the ray-finned fish *Polypterus*, which behaves as an outliner in some comparisons.

### Alternative Anatomical Interpretations

In building anatomical networks of appendicular skeletons, we included only bones and articulations whose presence we were confident about (“minimal networks”). However, 11 of the 34 appendages modelled had one or more articulations for which we were uncertain about their presence; mostly involving mediolateral contacts between bones in extinct taxa. We accounted for variations due to the presence/absence of these articulations by analysing network models that also included these uncertain articulations (“extended networks”). Regardless of minor changes in the values of some topological variables, the main results and interpretation for extended networks were the same as for minimal networks (see details in **Supplementary Results**). This indicated that our analysis could accommodate informed variations in network models, due to alternative interpretations of the anatomy of appendages, without nullifying the main results and conclusions of this study.

## DISCUSSION

The evolution of tetrapod limbs from fish fins occurred through a series of anatomical changes, including the loss/gain of girdle elements, acquisition of wrist/ankle joints, and the development of digits^5^. Here we demonstrated that digits had the greatest impact on the evolution of the anatomy of the appendicular skeleton. Topological variables discriminated between digit-bearing taxa and radial-bearing taxa, which occupied distinct regions of the morphospace (**Fig. 2**). Studies based on different appendicular skeletal traits also discriminated between digit-bearing taxa and radial-bearing taxa^30^. From a topological viewpoint, digits increased the number of bones (N) and articulations (K) of appendages, but in such a way (in contrast to carpal/tarsal topological arrangement of bones) that both the relative density of articulations (D) and the heterogeneity of number of articulations of bones (H) decreased. At the same time, phalanges articulating one-on-one proximodistally increased the overall path length (L) of appendages and the assortativity of bones (A). Finally, the formation of new digit modules increased the parcellation (P) of limbs compared to lobe-fins, making them more modular^22^. The origin of digits was concomitant with the parallel evolution of pectoral and pelvic limbs toward a region of the morphospace (**Fig. 2a**, top-right) where digit-related features predominated (high N, K, A, P; low D, H). Our results also showed a significant evolutionary shift in the topological anatomy for digit-bearing taxa (**Fig. 3**) that departed from the previous evolutionary regime observed in radial-bearing taxa, even in intermediate forms such as *Eusthenopteron*, *Gogonasus*, and *Tiktaalik*. These findings are not a surprise if one acknowledges the presence of digits as a morphological and evolutionary novelty^26, 31–33^, regardless of shared developmental histories or “deep-homologies” between digits and radials^34–38^. Novelties that increase the potential morphological morphospace provide an opportunity for greater diversification^39^. The evolutionary separation of digit-bearing taxa in the morphospace and in evolutionary estimations is in line with the idea of limbs as having evolved a truly novel anatomical reorganization^40^. This novelty has well-recognized functional implications for the origin of terrestrial locomotion, but the evidence for modularity may have more than developmental implications. By partitioning fins into more modular limbs there should have been more potential for localized functional specializations of bones, joints, muscles and more; including differential functions of pectoral and pelvic appendages^9^. Such potential would also arise from the modified degrees of freedom of the limb, transformed from somewhat homogeneous, flexible fins into fewer, more stiffened distal limb joints (i.e. in the autopodium). This is a logical speculation even though our analysis of bone topological networks is unable to account for details of the lepidotrichia.

The topological disparity of pectoral and pelvic appendages also decreased during the timespan between the origin of sarcopterygians and the origin of amniotes (**Fig. 4a**). A divergent pattern was observed for pectoral and pelvic appendages, when we considered all appendages studied (i.e., the complete sample). Whereas pectoral disparity decreased exponentially within this time interval, pelvic disparity only decreased linearly, until approximately the time of the earliest described tetrapod trackways (e.g., the Zachełmie tracks from the Middle Devonian^41^; similar in timing to other records pre-dating substantial body fossils^42,43^) and then it stabilized. This pattern, although based on a modest sample size as mandated by the existing fossil record, is congruent with an evolutionary stabilization of the disparity of pelvic anatomical organization near the origin of tetrapods, which did not occur for pectoral appendages until deeper within the tetrapod stem lineage. It is possible that this pattern is a result of few tetrapodomorph fish in our sample; when *Sauripterus*, *Gogonasus*, and *Tiktaalik* were removed from the pectoral analysis, we got the same pattern as for the pelvis. When the same taxa are included in the analyses, topological disparity decreased in parallel for pectoral and pelvic appendages, with both showing a shift in the decay rate coincident with the Zachełmie (and, approximately, other) tracks (**Fig. 4b**). If we consider these earliest tetrapod trackways part of the fins-to-limbs transition, our results would suggest that the transition indirectly decreased the morphological variation, which may have constrained the evolution of different topologies in limbs (see, for example, the morphospace occupation of limbs in **Fig. 2c**). This would agree with a dynamic compromise between possibly different functional demands in pectoral and pelvic appendages during the water-to-land transition^9^ and a shared developmental program constraining the evolvability of limbs^44^ (constraints that are still strong even in more deeply nested tetrapod lineages like primates^45^). Differing constraints and perhaps compromises have also been proposed to explain a decrease of disparity in the lower jaw of tetrapodomorphs across the water-to-land transition^46^. However, while functional trade-offs are likely for the feeding *vs.* locomotor systems of stem tetrapods, this de-coupling remains to be studied for the craniocervical region, which was likely more constrained in tetrapodomorphs by mechanical interactions between the pectoral appendage, axial column, and skull.

Previous studies have suggested the presence of bottlenecks in pectoral-pelvic anatomical similarity during the evolution of vertebrates^7,20,23–25^. One similarity bottleneck was proposed at the origin of tetrapods coincident with the fins-to-limbs transition^7,25^. This bottleneck was supported by data from general morphological features (e.g., shape and size similarities, presence/absence of homologous bones)^3^ and overall configuration and number of muscles^25^. Our disparity *vs.* time regression tests (**Fig. 5**) rejected the presence of this bottleneck for the eight topological variables measured for taxa across the fins-to-limbs transition. These new tests override our previous tentative support for this bottleneck from the analysis of the network-anatomy of extant taxa alone^22^. Our result, based on the network-anatomy or topological organization of the skeleton, differs of previous observations based on different skeletal and muscular features. One possible resolution to this apparent contradiction is that bones and muscles may have had a decoupled evolution during the fins-to-limbs transition, mirroring the idea that bones and muscles can respond differentially in time and magnitude to evolutionary pressures^47–49^. Lingering challenges for this question include the differing nature of data in bottleneck analyses (e.g. phylogenetic characters^3^; muscular attachments^25^) relative to this study, in addition to other complex traits not yet considered in such analyses, such as lepidotrichia.

Our study of the network-anatomy of appendages during the fins-to-limbs transition revealed an overall parallelism in the evolution of pectoral and pelvic appendages during this time, shaped greatly by the origin of digits. Digits were a morphological novelty that significantly changed the topological features of appendages, clearly discriminating limbs from fins and even from transitional forms. The presence of digits produced a directional evolutionary shift towards appendages that overall were less densely and heterogeneously connected, but more assortative and modular. Digits evolved in the context of a general decrease in topological disparity among pectoral as well as pelvic appendages, which may have had an impact on the subsequent evolution of tetrapods in terms of function, behaviour, and ecology.

## METHODS

### Anatomy of extinct taxa

We examined the skeletal anatomy of the pectoral and pelvic appendages in 11 extinct taxa: *Sauripterus taylory* Hall 1843; *Eusthenopteron foordi* Whiteaves 1881; *Gogonasus andrewsae* Long 1985; *Tiktaalik roseae* Daeschler, Shubin & Jenkins, 2006; *Acanthostega gunnari* Jarvik 1952; *Ichthyostega* sp. Säve-Söderbergh 1932; *Tulerpeton curtum* Lebedev 1984; *Balanerpeton woodi* Milner & Sequeira 1994; *Eryops megacephalus* Cope 1877; *Seymouria baylorensis* Broili 1904; and *Westlothiana lizziae* Smithson and Rolfe 1990. Our resources included museum collections, photographs and literature descriptions (see details in **Supplementary Materials**). These taxa were selected because they have articulated specimens with complete pectoral and/or pelvic appendages for examination and/or described in the literature. We only considered incomplete or disarticulated materials when a full, rigorous reconstruction of the appendage was available in the literature. Finally, we decided to exclude from the analysis those appendages for which the complete skeletal anatomy could not be confidently reconstructed due to a large number of missing elements, namely: pectoral appendages of *Acanthostega*, *Ichthyostega*, and *Westlothiana*; and pelvic appendages of *Sauripterus*, *Gogonasus*, and *Tiktaalik*.

### Anatomy of extant taxa

We examined the skeletal anatomy of pectoral and pelvic appendages in nine extant taxa. Six of them were recently described elsewhere^22^: *Polypterus senegalus* Cuvier 1829; *Latimeria chalumnae* Smith 1939; *Neoceratodus forsteri* Krefft 1870; *Ambystoma mexicanum* Shaw 1789; *Salamandra salamandra* Linnaeus 1758; and *Sphenodon punctatus* Gray 1842. In addition, we built network models for *Iguana iguana* Linnaeus 1758; *Didelphis virginiana* Kerr 1792; and *Mus musculus* Linnaeus 1758 (see details in **Supplementary Materials**). We selected these extant taxa because there were available dissection data and because they bracket the fins-to-limbs transition (i.e., rootward and crownward relative to Tetrapoda/Amniote).

### Network modelling

We built unweighted, undirected network models for the appendicular skeleton, where nodes coded for bones and links connecting nodes coded for physical articulation or contacts between two bones. Network models included the girdle and fin/limb skeleton. For the girdles, we considered all skeletal elements present or presumed as present: in pectoral girdles, these may include interclavicle, clavicle, supracleithrum, anocleithrum, cleithrum, and scapulocoracoid; in pelvic girdles these may include the hip bones fused (pelvis) or divided into two or three parts (ilium, pubis, and ischium). For fin and limb skeletons we considered all endochondral elements with a sufficient degree of ossification to be directly observed, as well as those elements for which there was enough indirect evidence (for example, an articular surface in another bone). We decided to exclude peripheral dermal elements, such as lepidotrichia and scales, from the fin network models for two main reasons. Firstly, it is often impossible to precisely identify their physical contacts to other elements in fossil taxa; secondly, their absence in digit-bearing taxa adds noise to the comparison of the skeletal topology between fins and limbs using network analysis.

We coded the articulations among bones following detailed descriptions of each taxon (see **Supplementary Materials**). When in doubt, we considered physical contiguity and adjacency as presence of articulation, which allowed us to code for contacts between bones in fossils that did not preserve details of the articular surface due to lack of preservation. Nevertheless, it was sometimes difficult to discern the presence/absence of a given contact between two bones in fossil taxa. We tackled this uncertainty at the modelling level and at the analysis level. At the modelling level, by building two types of networks for each appendage: a minimal network that includes the contacts with high certainty and an extended network that includes also potential, but more uncertain contacts (see **Supplementary Materials**). This assesses whether different criteria may affect the evolutionary patterns reported. At the analysis level, by performing a robustness tests of network parameter values under the assumption of random noise or sampling error (see below).

### Anatomical network analysis

We characterized the architecture of fins and limbs using eight topological variables (network parameters): number of nodes (N) and number of links (K), density of connections (D), mean clustering coefficient (C), mean path length (L), heterogeneity of connections (H), assortativity of connections (A), and parcellation (P). In short, parameters N and K are counts of the number of bones and physical contacts among bones, respectively. D measures the actual number of connections divided by the maximum number possible (it ranges from 0 to 1); D increases as new bones evolve if they form many new articulations, otherwise it decreases. C measures the average of the ratio of a node’s neighbours that connect among them (it ranges from 0 to 1); in the appendicular skeleton, triangular motifs can form by adding mediolateral articulations to the most commonly present proximodistal ones. L measures the average number of links required to travel between two nodes (minimum 1); L will increase, for example, by the presence of serial bones articulating one-on-one proximodistally. H measures the variability in the number of connections of nodes as the ratio between the standard deviation and the mean of the number of connections of all nodes in the network (minimum 0); appendages where each bone has a different number of articulations will have a high H, while if bones have the same number of articulations (i.e., forming a regular pattern) the appendage will have a low H. A quantifies the extent to which nodes with the same number of connections connect to each other (positive if nodes with the same number of connections connect to each other, negative otherwise); when this happens A is positive, whereas the inverse tendency means that A is negative. Finally, P measures the degree of modularity of the network (it ranges from 0 to 1); appendages with more network-modules and with bones evenly distributed among modules will have a high P. See the **Supplementary Materials** for further mathematical details. We measured topological variables in R^50^ using functions from the package *igraph*^51^.

### Parameter robustness

We tested the robustness of topological variables to potential errors in assessing the presence of bones and articulations by comparing the observed values to a randomly generated sample of 10,000 noisy networks for each anatomical network. We created noisy networks by randomly rewiring the links of the original network with a 0.05 probability, which results in introducing a 5% artificially generated error. Then, we compared the observed values of empirical networks to the sample of noisy networks. In each case, we tested the null hypothesis that observed values are equal to the sample mean. We rejected the null hypothesis with α = 0.05 if the observed value is in the 5% end of the distribution of simulated values (**Supplementary Table 2**; “TRUE”, cannot reject H_0_; “FALSE”, reject H0 with α = 0.05). We tested a total of 272 values (34 networks x 8 parameters): 268 fell within the confidence intervals and scored “TRUE” in the test. The exceptions were for *Neoceratodus* pectoral path length, *Neoceratodus* pelvic clustering coefficient and path length, and *Didelphis* pelvic parcellation. Because the anatomy of *Neoceratodus* and *Didelphis* derived from our own dissections, these few cases of rejection of the null hypothesis for these parameters can be attributed to the difficulty for a random-noise process to produce realistic dissection errors (for example, by coding the femur as not articulating with the pelvis).

### Phylogenetic relationships

We assembled a phylogenetic tree for our study taxa according to the approximate majority view in published phylogenies^52–54^. We calibrated the tree branches using the ‘equal’ method defined by Lloyd^55,56^ to adjust zero-length and internal branches, as implemented in the package *paleotree*^57^ for R. Temporal ranges of taxa (first and last appearance date in Ma) follow those of the Paleobiological Database (www.paleodb.org) and TimeTree (www.timetree.org) (**Supplementary Tables 4 and 5**). We constrained tree calibration by assigning minimum dates for known internal nodes (clades or splits) based on molecular inferences and fossil dates from the literature (*op. cit.*). The exact dates for first and last appearance of taxa and for internal nodes are available within source code attached. Note that when required by the analysis, the main tree was pruned to only include those taxa of interest.

### Analysis of topological variation

We ran a principal component analysis (PCA) of topological variables by a singular value decomposition of the centred and scaled measures using the function *prcomp* in the R build-in package *stats*^50^. We used PCA components to test whether the anatomical organization of the appendicular skeleton differed (1) between fins (without digits) and limbs (with digits), (2) between pectoral and pelvic appendages, and (3) between extinct and extant taxa. We performed a permutational multivariate analysis of variance (PERMANOVA) over 10,000 permutations using the function *adonis* in the R package *vegan*^58^. PERMANOVA used a permutation test with pseudo-F ratios on the Euclidean distances of the matrix of PCA components to test the null hypothesis that the centroids and dispersion were equivalent for each group comparison. Rejection of the null hypothesis meant that the network topology differed between the groups compared.

### Evolutionary modelling

We estimated the occurrence of evolutionary shifts in the topological organization of appendages in our phylogenetic tree using a SURFACE^59^ analysis of the first two PC components for pectoral and pelvic appendages, independently. SURFACE estimates change of evolutionary regimes—in the strength (α) and rate (σ) of evolution and in the optimal mean (θ)— from multivariate data and a non-ultrametric tree. SURFACE uses an Ornstein-Uhlenbeck (OU) stabilizing selection model of evolution, which allows changes in the rate of evolution and optimal means of variables. If present, this method identifies homoplasy: two clades with the same regime. Given the small sample of appendages in our comparisons, we deemed it necessary to calculate the power of the SURFACE analysis as an indicator of reliability in the accuracy of estimated patterns. Because in OU models power is dependent by strength, rate, and optimal mean combined, effect-size measures offer a better prediction of power than sample size^60^. We calculated the power of the estimated regimes using the Signal-to-Noise Ratio (SNR), which is defined as 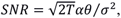 where T is the total depth of the phylogeny. High power can be inferred when SNR >>1. To further validate the estimated regimes, the output of SURFACE was then fitted to alternative evolutionary models: a 1-rate Brownian Motion (BM); a multi-rate BM; an OU with fixed strength, rate, and mean; an OU with fixed strength and rate, and multi-mean; and an OU with fixed strength, and multi-rate and multi-mean. Fitted models were compared using Akaike Information Criteria.

Finally, we statistically tested the resulting evolutionary models of evolution with a phylogenetic MANOVA to confirm that the clades identified had a different regime corresponding to different groups with different means. The combination of estimation, fitting, and testing allowed us to build confidence that the evolutionary patterns found were reliable if they converged on the same result. Evolutionary modelling was carried out in R using functions from the packages *surface*^59^, *mvMORPH*^61^, and *geiger*^62^ for the estimation, fitting, and testing, respectively.

### Disparity through time (DTT)

To examine how topological disparity changed over time, we performed a disparity through time (DTT) analysis on pectoral and pelvic appendages, separately. Following a previous study on mammalian neck anatomy^63^, first we obtained the co-variation of topological variables by performing independent PCAs for pectoral and pelvic networks. Next, we calculated the mean subclade disparity on the PC scores using the function *dtt* in the R package geiger^62^. The higher the disparity, the higher the variance within subclades (i.e., lower conservation) and the lower the variance between subclades^64,65^. We tested the statistical significance of the observed disparity with a randomization inference test with 10,000 simulations under a Brownian motion evolution on our phylogeny. Probability values were calculated empirically at each subclade time and combined using Edgington’s method^66^ as implemented in the R package *metap*^67^. Function *dtt* also calculated the morphological disparity index, which quantified the overall difference in relative disparity of a clade compared to that expected under the null Brownian motion model. For reference, DTT analyses were also performed on the subsample of taxa for which we have both pectoral and pelvic appendages.

### Pectoral-pelvic similarity bottleneck

We tested the hypothesis of the existence of a pectoral-pelvic similarity bottleneck at the origin of Tetrapoda for each topological variable independently. For practical purposes we used pectoral-pelvic disparity (lower disparity means greater similarity). Pectoral-pelvic disparity was calculated as the absolute residuals of pectoral and pelvic values on the identity line (or 1:1 line, a line with intercept=0 and slope=1), so that identical pelvic and pectoral appendages—maximal similarity—had a value of zero disparity. According to previous formulations of the bottleneck hypothesis for tetrapods, taxa before and after the split of tetrapods would have a greater pectoral-pelvic disparity (lower similarity) than taxa closer to the origin of tetrapods^7,22,25^. Thus, the farther we go in time from this event the greater the expected pectoral-pelvic disparity should be. To test this prediction, we performed a phylogenetic generalized least square regression (PGLS) of the absolute pectoral-pelvic residuals on the 1:1 line against the time from the Tetrapoda branch of taxa having both appendages in the sample (t_0_=370.4 My). We used PGLS to test against the null hypothesis of a slope=0, meaning no difference in pectoral-pelvic disparity through time. For our prediction to hold, pGLS needed to show a statistically significant positive regression slope. PGLS was computed in R using a standard generalized least-square with an *a priori* correlation structure derived from the phylogenetic tree using the function *corPagel* of the package *ape*^68^.

## ACKNOWLEDGEMENTS

We thank Jennifer Clack, Timothy Smithson, Michael Coates, and Roger Benson for their help interpreting skeletal anatomy in fossil specimens. John Hutchinson’s team at RVC is thanked for their input on early drafts. We thank Jeroen Smaers for his teaching and insight on phylogenetic analyses. We also thank Jessica Cundiff (Museum of Comparative Zoology at Harvard) and Emma Bernard (Natural History Museum of London) for their assistance in accessing specimens. This project is funded by the European Union’s Horizon 2020 research and innovation program under the Marie Skłodowska-Curie grant agreement No 654155.

## Reporting Summary

Further information on experimental design is available in the Nature Research Reporting Summary linked to this article.

## Data Availability

The data that support the findings of this study are available from Figshare at https://figshare.com/s/4c3a1dd62d1f55728a0e.

## Code Availability

The code that support the findings of this study are available from Figshare at https://figshare.com/s/4c3a1dd62d1f55728a0e.

## Materials & Correspondence

Borja Esteve-Altava

## Author Contributions

BE-A, SEP, JRH: Built and revised the adjacency matrices for extinct taxa

BE-A, JLM, PJ, JRH, RD: Built and revised the adjacency matrices for extant taxa

BE-A: designed the study, analysed the networks, and wrote the manuscript.

All authors discussed the results and revised the manuscript.

